# Conserved RNA structural elements mediate WDR5 recognition of the long non-coding RNA UMLILO

**DOI:** 10.64898/2026.11.07.765059

**Authors:** Amy Crawford, Alfredo Vidal-Ceballos, Alisha N. Jones

**Affiliations:** Department of Chemistry, New York University, New York, New York, USA

**Keywords:** UMLILO, long noncoding RNA, protein, WDR5, structure, innate immunity, biomolecular condensates

## Abstract

The long non-coding RNA (lncRNA), UMLILO (upstream master lncRNA of the inflammatory chemokine locus), regulates the epigenetic programming of innate immune genes. Although its basic mechanism was revealed in cells, the molecular basis by which UMLILO interacts with its partner, WD repeat-containing protein 5 (WDR5), remained unknown. To understand how UMLILO interacts with WDR5, we determined the secondary structure of UMLILO by chemical probing and performed binding assays, uncovering the multivalent nature of their interaction, involving multiple RNA stem-loops. Conservational analysis revealed a stem-loop structurally conserved across mammals. Furthermore, we found that UMLILO engages multiple sites on the surface of WDR5 as revealed by NMR chemical shift perturbation analysis, and that full-length UMLILO increases WDR5 recruitment into *in vitro* condensates. Our report provides a detailed characterization of the molecular basis of WDR5 recognition by an RNA and establishes a new paradigm for the role of IPLs in enhancing gene expression.

## Introduction

Innate immune cells can undergo epigenetic reprogramming to enhance their response to infection. This phenomenon, known as trained immunity, is characterized by increased histone H3 lysine 4 trimethylation (H3K4me3) at immune gene promoters^1,2^. Immune gene-priming long non-coding RNAs (IPLs) are required for H3K4me3 accumulation at the promoters of innate immune genes^3^. By regulating IPL expression, immune cells can fine-tune inflammatory responses to respond quickly to invading pathogens while avoiding excessive inflammation. IPLs are transcribed near the promoters of target genes, where they act in *cis* to direct chromatin-modifying proteins to nearby promoters. This ability is facilitated by nuclear architecture, in which co-regulated genes are brought near the site of IPL transcription through the arrangement of chromosomal loops into a nuclear compartment called a topologically associating domain (TAD). Here, the IPL recruits the WD repeat-containing protein 5 (WDR5)/Mixed Lineage Leukemia 1 (MLL1) complex, which catalyzes H3K4 trimethylation (**Fig. 1A**).

**Figure 1.**
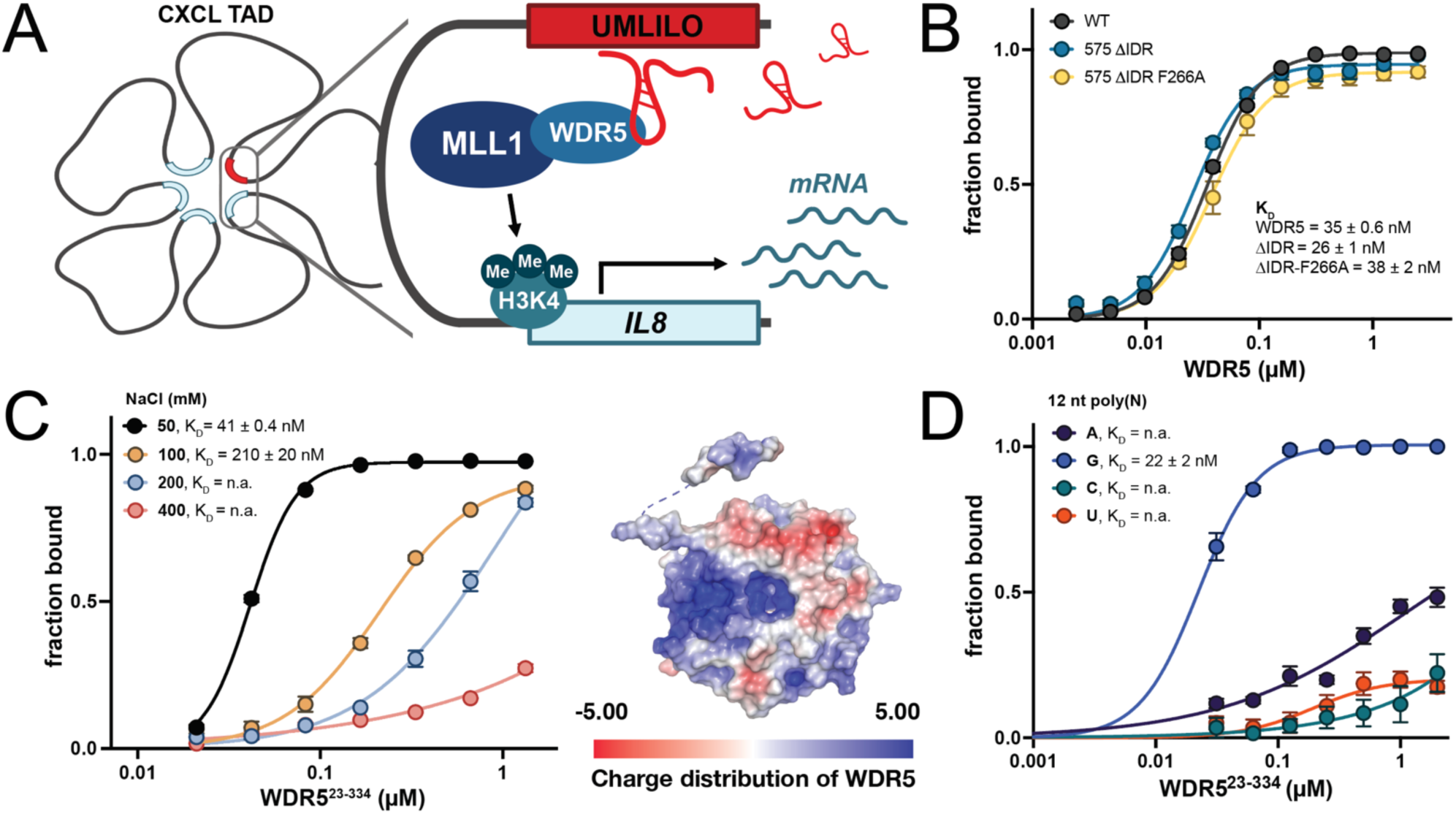
UMLILO binds WDR5 with high affinity in an electrostatic-dependent manner. **A.** When UMLILO transcription is induced, UMLILO acts *in cis* to bind WDR5 to recruit the MLL1/WDR5 complex to the CXCL chemokine TAD, where MLL1 primes the immune genes for upregulation by catalyzing trimethylation of H3K4. Adapted from Fanucchi et al., 2019. **B.** Binding curves of WDR5 constructs and UMLILO. **C.** Binding curve of WDR5 and UMLILO under increasing salt concentrations (left), charge distribution mapped on the surface of WDR5 (right), showing the positively charged region around the WBM site. PDB: 2GNQ. **D.** Binding curves of polynucleotide RNAs to WDR5. UMLILO = Upstream Master LncRNA of the Inflammatory chemokine Locus, WDR5 = WD repeat-containing protein 5, MLL1 = Mixed Lineage Leukemia 1, TAD = Topologically Associating Domain, H3K4 = Histone H3, Lysine 4, WBM = WDR5 Binding Motif. For all binding assays error bars represent s.e.m., n = 3 technical replicates.

The IPL Upstream Master LncRNA of the Inflammatory chemokine LOcus (UMLILO) is a 575-nucleotide (nt) transcript that is responsible for training *IL8, CXCL1, CXCL2,* and *CXCL3* chemokine genes within the CXCL TAD in humans^3^. These chemokines induce neutrophil chemotaxis to the site of infection and are among the first lines of defense in mammalian immune responses. UMLILO facilitates this process by interacting directly with WDR5, a highly conserved scaffolding protein composed of seven β-sheet propellers with two hydrophobic binding pockets on either side connected by a hydrophobic central channel^4^. Several residues have been proposed to be important for this interaction, including a surface-exposed phenylalanine (F266) that, when mutated to an alanine, completely abolished WDR5’s ability to interact with RNA in cells^5^. Following WDR5 binding, UMLILO acts as a guide to recruit the Mixed Lineage Leukemia 1 (MLL1) methylation complex to prime the *CXCL* gene cluster.

Though studies are sparse, WDR5 has been found to partition into liquid-liquid phase-separated (LLPS) condensates both *in vitro* and in cells. This has revealed, for example, that i) LLPS increases the methylation activity of the MLL1/WDR5 complex 60-fold^6^, ii) several proteins co-localize to biomolecular condensates with WDR5, including NANOG and MYC, to potentially activate gene expression^7,8^, and iii) RNA influences the formation of WDR5 condensates^8^. Here, fragments of another well-known WDR5-binding lncRNA, HOTTIP, were found to increase WDR5’s partitioning into condensates. These results are in line with the growing evidence that phase separation plays an important role in organizing and regulating modifying protein complexes at gene promoters, and that TADs can contain non-coding RNA-nucleated condensates that modulate TAD structure and drive gene expression^9–12^.

WDR5’s structural properties have been well characterized, and it is of high therapeutic interest due to its involvement in multiple oncogenic pathways^13–16^. WDR5 has two known binding pockets: (1) the WDR5-interacting motif (WIN) site, which mediates chromatin and MLL1 interactions; and (2) the WDR5 binding motif (WBM) site, which binds MYC and the RbBP5 subunit of the MLL1/WDR5 complex; both of which have been therapeutically targeted^17–20^. The multifunctionality of WDR5, however, makes it a challenging target for the development of tools for specific control of epigenetic modulation^21–23^. An alternative approach would be to target WDR5–RNA complexes. Notably, WDR5 binds to over 100 long non-coding RNAs, such as HOTTIP, ANRIL, NeST, FOXD3-AS1, and MERRICAL, transcripts that are important for the localization of WDR5 to chromatin^5,24–29^. Many of these RNAs, including HOTTIP and IPLs, are transcribed under strict spatial and temporal control, thereby restricting WDR5-mediated upregulation to the nuclear compartment where the RNA is transcribed^3,29^. Interfering with these *cis-*acting lncRNA– WDR5 interactions using a small molecule or biologic would achieve specific gene suppression or activation, making them a valuable target. Designing such therapeutic agents requires information regarding the structural mechanisms underlying WDR5–RNA interactions and functions^30^.

Targeting these WDR5–RNA interfaces requires the investigation of the atomistic and structural details that underlie these interactions. Here, we combine biochemical and biophysical approaches to characterize the interaction between WDR5 and UMLILO. Using binding assays, we find that UMLILO binds to WDR5 with nanomolar binding affinity, an interaction driven by electrostatics. RNA structural probing experiments and conservation analysis reveal that UMLILO is intricately structured and harbors conserved stem-loop motifs responsible for WDR5 binding. NMR titrations reveal two β-sheet propellers that primarily serve as the binding site for UMLILO, and that RNA binding to WDR5 perturbs residues of the WIN site essential for MLL1 binding. Finally, we show that UMLILO promotes WDR5 condensate formation, and properties of WDR5 condensates are influenced by RNA length and structure, providing *in vitro* evidence that the size and structure of UMLILO are both important for its function in recruiting WDR5. Overall, this work provides an in-depth biochemical and biophysical analysis of WDR5–RNA binding and is the first to demonstrate that specific secondary structural elements of an IPL bind WDR5. This work contributes to our overall understanding of how WDR5 interacts with lncRNAs to perform vital cellular functions and paves the way for identifying and designing inhibitors that block the WDR5–UMLILO interaction.

## Results

### WDR5 binds to UMLILO with nanomolar binding affinity

UMLILO and WDR5 directly interact in the nucleus, as shown by cross-linking immunoprecipitation, to recruit the full WDR5/MLL1 complex to *CXCL* gene promoters (**Fig. 1A**)^3^. Filter dot blot binding assays of UMLILO and WDR5 were performed to validate complex formation *in vitro* and to further quantify and characterize this interaction. WDR5 bound UMLILO with low nanomolar affinity (K_D_ = 35 ± 0.6 nM), which was improved slightly by truncating the intrinsically disordered region (ΔIDR = WDR5^23–334^, K_D_ = 26 ± 1 nM; **Fig. 1B, Supplemental Fig. 1A**). Their interaction was cooperative as demonstrated by a Hill coefficient of 1.9 ± 0.05, consistent with multiple binding events or conformational changes induced by protein binding.

It has previously been shown that mutating phenylalanine 266 disrupts WDR5 interactions with lncRNAs *in vivo*. Despite widespread use of this F266A mutant for studying lncRNA–WDR5 interactions^5,28,31^, this mutation had minimal effect on the WDR5–UMLILO interaction *in vitro* (ΔIDR-F266A K_D_ = 38 ± 2 nM; **Fig. 1B, Supplemental Fig. 1A**). This may be due to differences between binding *in vitro* and long-term cellular stability of the F266A mutant. To further investigate the effect of F266A on WDR5 structural stability, we performed circular dichroism (CD) and thermal denaturation assays. We found that, although the proteins are similarly folded, the F266A mutant had a lower melting temperature relative to wild-type WDR5 (40 ± 0.2°C and 56 ± 0.4°C, respectively; **Supplemental Fig. 1B&C**). The F266 residue is located within a hydrophobic pocket of WDR5, where the aromatic side chain may be essential for local structural stability. AlphaFold models of WDR5 mutants support this hypothesis. When residues in the WBM pocket implicated in lncRNA binding were each substituted with an alanine, only the F266A model contained a potentially destabilizing channel between β-propellers (**Supplemental Fig. 1D**)^32^. Taken together, this evidence suggests that F266 is unlikely to be directly responsible for the interaction between WDR5 and UMLILO and may instead be essential for structural stability.

The binding affinity of the wild-type protein was sensitive to changes in ionic strength and decreased fivefold when the sodium concentration increased from 50 mM to 100 mM (K_D_ = 41 ± 0.4 nM and 210 ± 20 nM, respectively), and was nearly abolished at 400 mM NaCl (K_D_ > 2.5 µM, above the highest protein concentration used in the assay; **Fig. 1C, Supplemental Fig. 1E**). The surface of WDR5 features several charged clefts, including a positively charged region that contains the WDR5 binding motif (WBM) site (**Fig. 1C**). The dependence of RNA binding on the ionic environment suggests WDR5 interacts near the WBM site *via* electrostatic interactions with the negatively charged RNA backbone. The interaction between UMLILO and WDR5 being heavily dependent on electrostatics could imply that WDR5 binds RNA promiscuously. To this end, we performed binding assays and found that WDR5 in fact had minimal binding to poly(A), (U), and (C) RNAs (**Fig. 1D, Supplemental Fig. 1F**). WDR5 did bind to poly(G) RNA, indicating that it may bind G-rich RNAs or G-quadruplexes elsewhere in the nucleus (K_D_ = 22 ± 2 nM, **Fig. 1D**).

### UMLILO adopts an intricate multi-domain secondary structure

The 575-nt canonical transcript was chemically probed with 1-methyl-7-nitroisatoic anhydride (1M7) and dimethyl sulfate (DMS) and revealed to have an intricate secondary structure (**Fig. 2A&B, Supplemental Fig. 2A**). The reactivity was in good agreement with the predicted 2D structures, with single-stranded regions typically showing greater reactivity than base-paired nucleotides. UMLILO contains two domains connected via a multiway junction. Domain I (nts 69–322) contains four main stem-loop structures, and domain II (nts 343–546) contains two stem-loops connected by another multiway junction.

**Figure 2.**
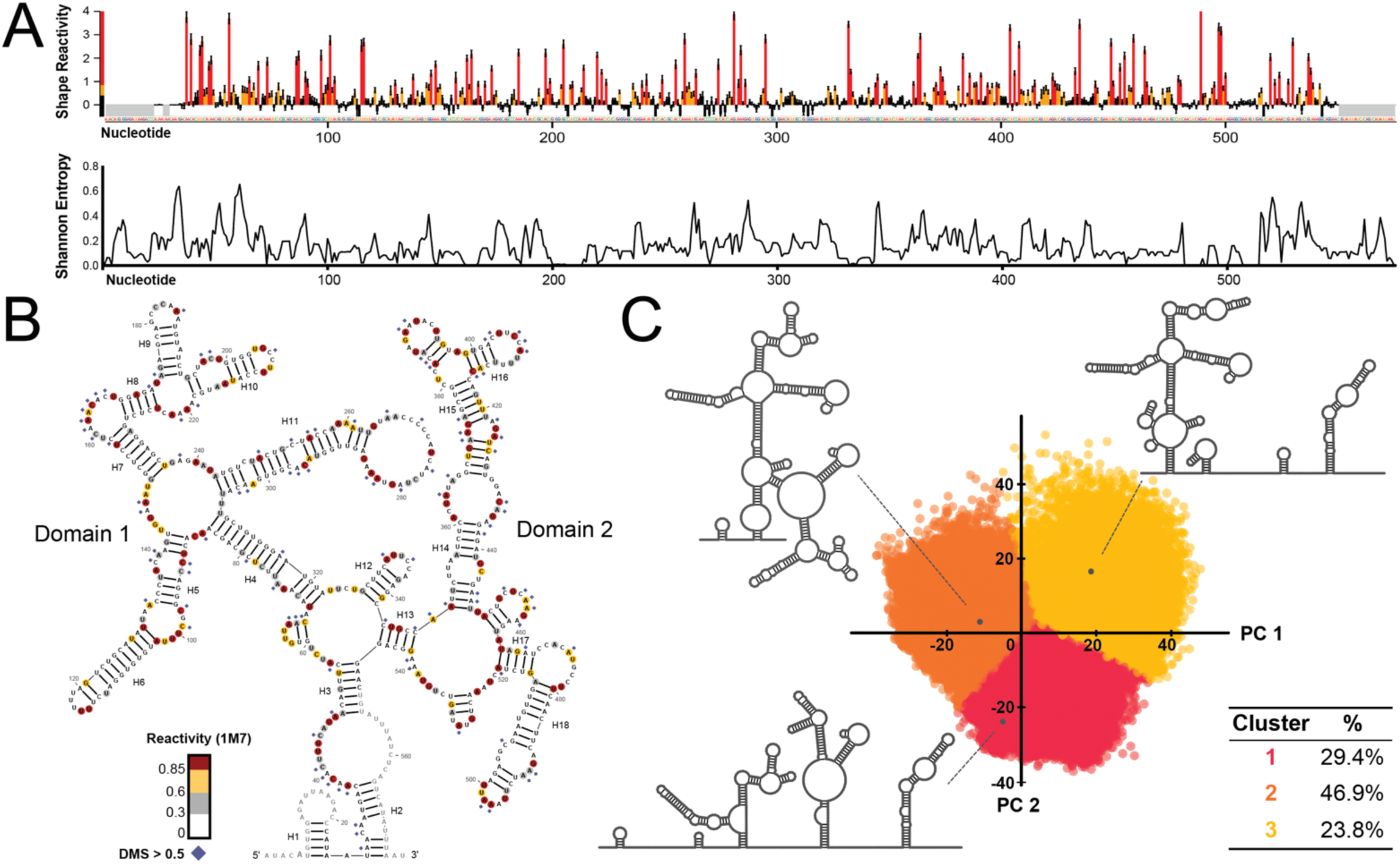
UMLILO is conformationally dynamic. **A.** SHAPE reactivity profile and plot of Shannon entropies of UMLILO probed with 1M7. **B.** Secondary structure of UMLILO informed by SHAPE data. **C.** DaVinci analysis of UMLILO probed with 5-NIA with representative alternative conformations. 1M7 = 1-methyl-7-nitroisatoic anhydride, DMS = dimethyl sulfate, 5-NIA = 5-nitroisatoic anhydride, PC = Principal Component.

The regions with high Shannon entropies along the length of the transcript suggest that UMLILO may adopt multiple conformations; in agreement with this, there are regions where the single-stranded nucleotides (e.g., in the apical loops of H11) lack the expected reactivity, a common indicator of a structurally dynamic RNA (**Fig. 2A**)^33,34^. DaVinci deconvolution analysis^35^ of the mutational profiles of UMLILO shows that it is indeed conformationally dynamic, with three broad clusters being identified, represented at 29.4% (14,457 structures), 46.9% (23,044 structures), and 23.8% (11,683 structures) (**Fig. 2C**). The dynamic regions occur particularly amongst nucleotides 140–300 of domain I and 360–440 in domain II.

### Identification and characterization of specific WDR5 binding regions on UMLILO

Chemical probing (both 1M7 and DMS) was performed on the UMLILO–WDR5 RNA–protein complex to identify protein-binding regions. In these experiments, regions protected by the protein will be shielded from the chemical probe, while regions where structural changes were induced may show increased (**Fig. 3A**, green bars) or decreased reactivity (**Fig. 3A**, purple bars).

**Figure 3.**
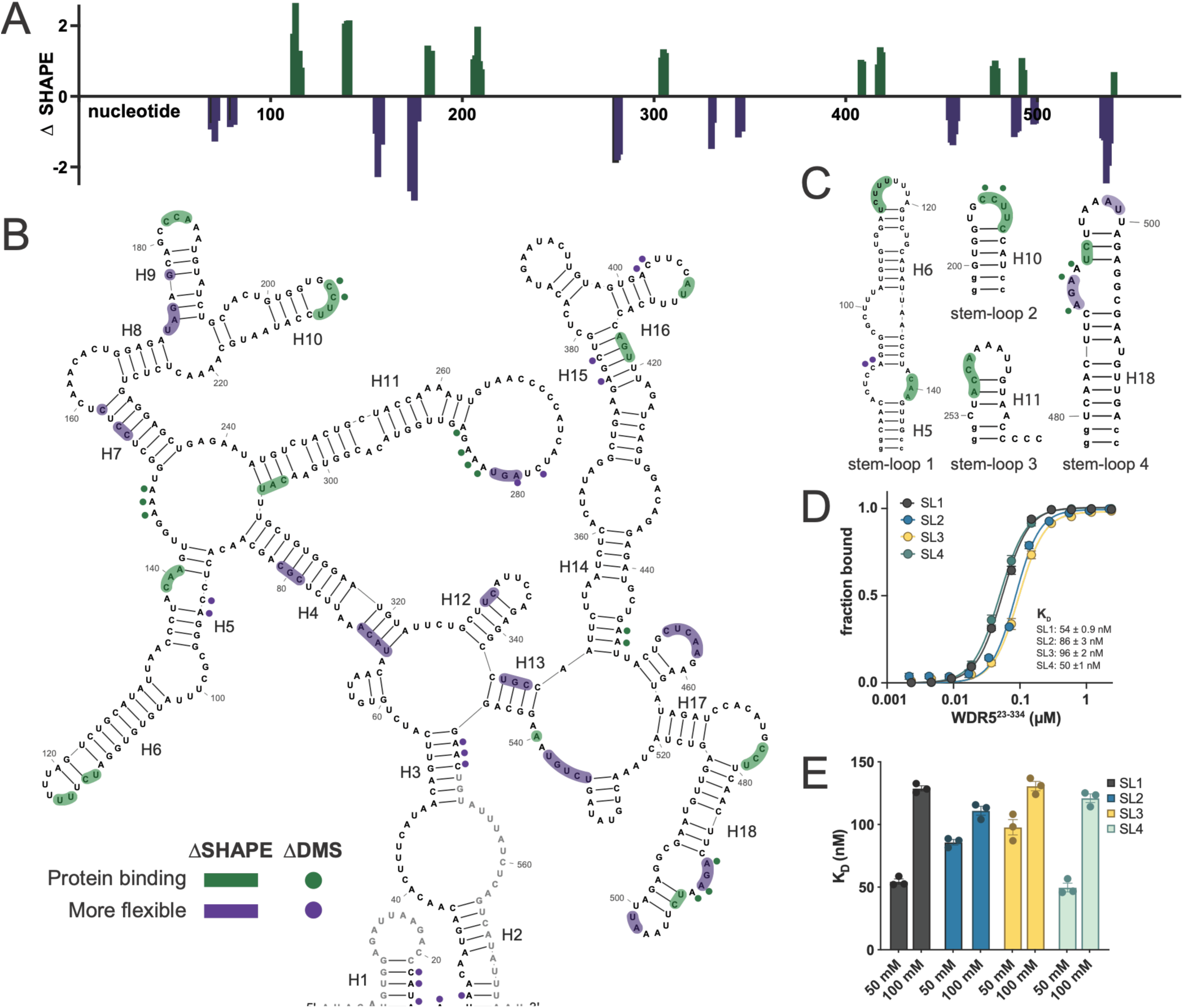
WDR5 binds multiple UMLILO structural motifs. **A.** Plot of changes in 1M7 reactivity of UMLILO in the presence of WDR5. Positive ΔSHAPE values (green) indicate decreased nucleotide flexibility consistent with WDR5 binding, while negative values (purple) correspond with increased flexibility. **B.** Secondary structure of UMLILO with highlighted regions showing WDR5 binding sites (green) and structural changes induced by protein binding (purple). Nucleotides with changes in reactivity to 1M7 are highlighted, changes in reactivity to DMS are represented by dots. **C.** WDR5–binding stem-loops identified through deltaSHAPE analysis. The stem-loop 3 structure and indicated WDR5-binding region are based on an alternative conformation and differences in reactivity to 5-NIA. Lowercase letters indicate nucleotides added for *in vitro* transcription of the stem-loop constructs. **D.** Binding curves for stem-loops and WDR5. Error bars represent s.e.m., n = 3. **E.** Affinity of each stem-loop for WDR5 under differing ionic conditions. Data points represent K_D_ from three replicates, error bars represent s.e.m. SL = stem-loop.

Overall, changes in 1M7 and DMS reactivity induced by the presence of WDR5 occurred in similar and sometimes complementary local regions of the RNA (**Fig. 3A&B, Supplemental Fig. 3A–C**). For example, nucleotides 206–209 (in H10) and nucleotides 487–493 (in H18) experienced reactivity changes for both probes. H5 reactivity changes were complementary, where DMS modified the 5′ half of the internal loop while 1M7 showed reactivity changes on the 3′ side. These results suggest that WDR5 engages not only with the ribose-phosphate backbone but also with the nucleobases. Notably, nearly all changes in the reactivity profiles were enriched among single-stranded, pyrimidine-rich nucleotides (e.g., helices H6, H9, H10, H16, H18). In addition to regions of decreased reactivity associated with protein binding, several regions became more flexible, indicating that WDR5 may indeed induce conformational changes in UMLILO.

To validate the binding sites, four hairpins that showed strong differences in reactivity in the presence of WDR5 (stem-loop (SL) 1 = nts 84–145, SL2 = 200–213, SL3 = 253–272, and SL4 = 480–515) were individually transcribed, and their binding with WDR5 was assessed (**Fig. 3C**). Of the stem-loops, the longer, more structured SL1 and SL4 bound WDR5 with the highest affinity (SL1 = 54 ± 0.9 nM, SL4 = 50 ± 1 nM; **Fig. 3D, Supplemental Fig. 3D**). SL2 and SL3 bound with lower affinity (SL2 = 86 ± 3 nM, SL3 = 96 ± 2 nM); notably, SL2 and SL3 fall within regions identified as conformationally dynamic and may bind to WDR5 more effectively when they sample other structured states.

We further investigated the binding of stem-loops to WDR5 in the presence of increased salt to determine whether their binding mechanisms differ from those of full-length RNA. The binding of SLs 1 and 4 was affected by increasing ionic strength, supporting the notion that recognition of stem-loops is dependent on electrostatic interactions that impart structural selectivity to binding (**SL1**: 50 mM NaCl = 54 ± 0.9 nM, 100 mM NaCl = 128 ± 2 nM; **SL4:** 50 mM NaCl = 50 ±1 nM, 100 mM NaCl = 120 ± 4 nM; **Fig. 3E, Supplemental Fig. 3D**). In contrast, SLs 2 and 3, which are much smaller and occur in conformationally dynamic regions, showed little change in binding affinity with increased NaCl (**SL2:** 50 mM NaCl = 86 ± 3 nM, 100 mM NaCl = 110 ± 3 nM; **SL3:** 50 mM NaCl = 96 ± 2 nM, 100 mM NaCl = 130 ± 4 nM). We propose that SL1 and SL4 contribute, in part, to the high affinity electrostatic nature of the WDR5:UMLILO interaction, while shorter hairpin in dynamic regions, such as SL2 and SL3, confer tolerance to the ionic environment.

### Functional conservation of UMLILO

Conservation of sequence and structure is a strong indicator of biologically functional motifs. We carried out phylogenetic analysis and multiple sequence alignment of UMLILO across 43 species and found that, although the overall sequence is only moderately conserved (pairwise identity of 64%), many regions showed high sequence conservation (**Fig. 4A, Supplemental Fig. 4A–C**). Exon 3 contains the most conserved nucleotides, many of which are contained within a transposable LINE element. Interestingly, the two small, flexible WDR5-binding stem-loops are located within this conserved region, supporting their potential functional conservation (**Fig. 4A**).

**Figure 4.**
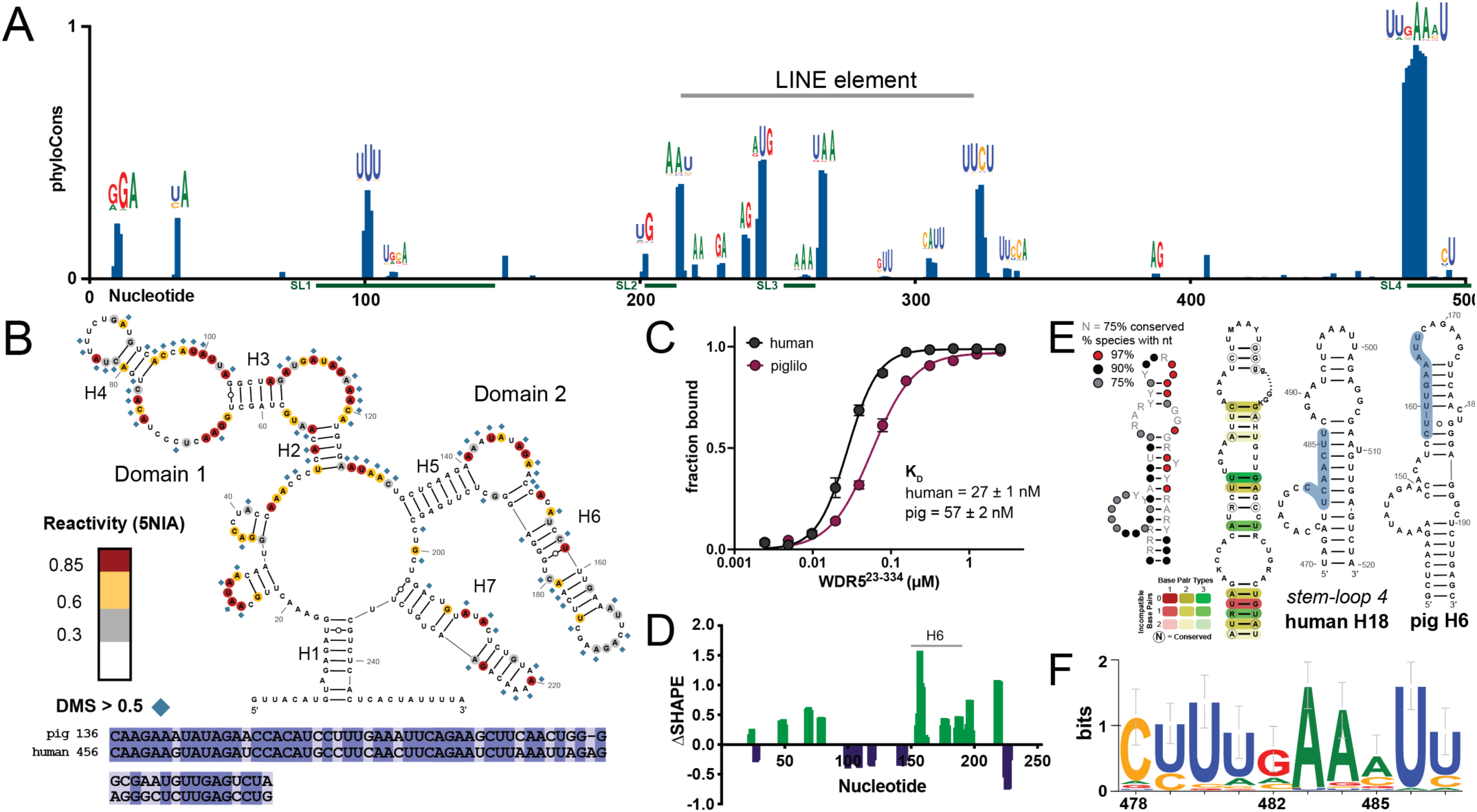
UMLILO is conserved across divergent species in structure and sequence. **A.** PhyloCons score for UMLILO from Hiller Lab 470 mammals sequence alignment dataset. Sequence logos from multiple sequence alignment are shown above conserved nucleotides. Regions corresponding to WDR5-binding SLs are indicated in green. **B.** Secondary structure of UMLILO in pigs shown above region of human and pig UMLILO sequence alignment with the highest similarity. **C.** Binding curve of pig and human UMLILO with WDR5. Error bars represent s.e.m., n = 3. **D.** deltaSHAPE plot for pig UMLILO at a 2:1 ratio WDR5:RNA. Positive values (green) indicate potential binding regions, while negative values (purple) indicate regions with increased flexibility. **E.** Secondary structure of the most conserved element: per-nucleotide conservation from R-scape analysis and RNAalifold consensus structure from LocARNA analysis are shown alongside human SL4 and pig helix 6. The most conserved sequence motif is highlighted in blue. **F**. Sequence logo of the most highly conserved motif from the multiple sequence alignment of UMLILO. LINE = Long Interspersed Nuclear Element, SL = stem-loop, R = A or G, Y = C or U, K = G or U, M = A or C.

We sought to validate these bioinformatic findings by determining the secondary structure of the pig homolog of UMLILO and assessing its binding to WDR5. Pig UMLILO is much shorter and aligns almost exclusively with the sequence corresponding to domain II of human UMLILO (**Supplemental Fig. 4D**) yet retains the ability to prime *IL8* chemokine genes^3^. Pig UMLILO was found to contain two domains: a more flexible domain (corresponding to H14–H15 in human UMLILO), and a much less reactive second domain containing a hairpin with high sequence and structural similarity to human UMLILO SL4 (**Fig. 4B, Supplemental Fig. 4E**). Despite sharing only 34% sequence identity, the pig homolog of UMLILO binds to WDR5 with relatively high affinity, on the order of human SL4 (pig UMLILO K_D_ = 57.1 ± 2 nM; **Fig. 4C, Supplemental Fig. 4F**).

Chemical probing in the presence of WDR5 revealed that hairpin H6, structurally similar to human SL4, is a WDR5 binding site (**Fig. 4D, Supplemental Fig. 4G**). The structure conservation of this hairpin is further supported by bioinformatic tools that perform structure-based alignments (**Fig. 4E, Supplemental Fig. 4H&I**). R-Scape analysis showed that nucleotides corresponding to SL4 are present in most species, with many of them fully conserved. LocARNA alignment of UMLILO domain II from twelve diverse species revealed a consensus structure containing a stem-loop that closely resembles SL4, showing structural conservation of base pairs. The sequence and structure conservation of SL4, as well as the shared ability of human and pig UMLILO to bind WDR5, supports the conclusion that WDR5 recognizes a specific hairpin structure.

Looking closely at conserved regions and WDR5-binding sites in human and pig UMLILO, we identified a pyrimidine-rich motif, 5′ <u>YUUVAANU</u> 3′ (N = any nucleotide; V = A, C, or G; Y = C or U) that is highly conserved across species (**Fig. 4A&F**). This conserved sequence is found twice in SL4 and pig H6, as well as in other lncRNAs that interact with WDR5 for epigenetic priming (**Fig. 4E**, **Supplemental Table 1**). Future eCLIP data and motif analysis will be valuable for determining whether this is a general WDR5 RNA-binding motif.

### RNA interacts with multiple surfaces of WDR5

NMR titration experiments were performed with WDR5 and UMLILO SL4 to determine the impact of WDR5 binding on the RNA structure and to identify amino acid residues on the WDR5-binding interface. Following the titration of RNA into protein, we observed that the imino resonances (11–15 ppm), which correspond to base-paired G’s and U’s of SL4, were broadly maintained, with expected line broadening (**Fig. 5A**). This indicates that WDR5 binds, but does not completely unwind, SL4, which is in agreement with the modest reactivity changes we observed in chemical probing experiments.

**Figure 5.**
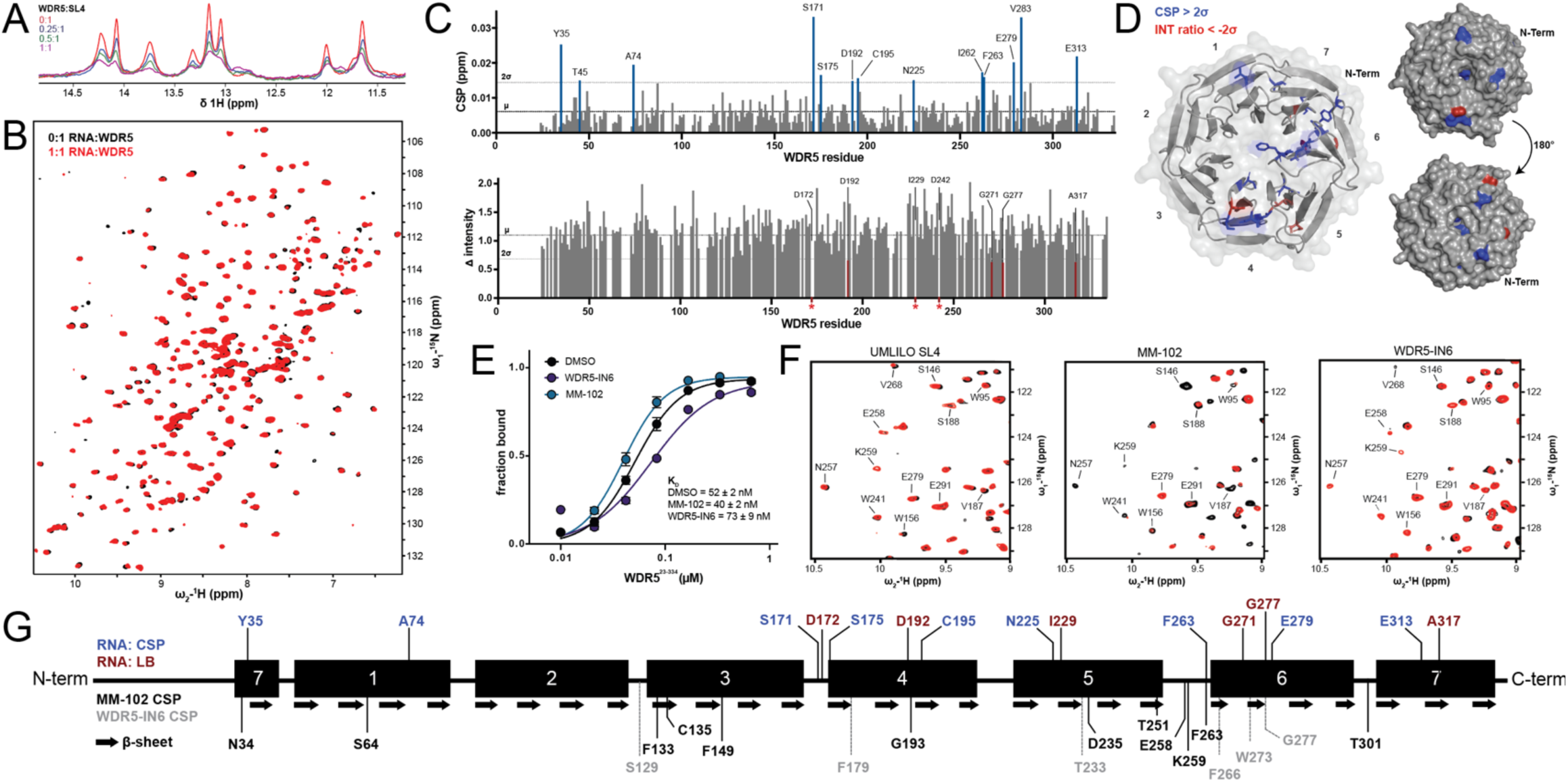
RNA and small-molecule inhibitors bind moderately distinct regions of WDR5. **A.** Overlay of 1D NMR spectra of UMLILO SL4 imino signals at different WDR5 titration points. **B.** ^1^H-^15^N TROSY NMR spectra of ^15^N-labeled WDR5^23–334^ alone (black) and in the presence of an equimolar amount of UMLILO SL4 (red). **C.** Plots of CSPs and changes in NMR peak intensity nduced upon addition of SL4 RNA to WDR5. Fully broadened peaks are represented by red asterisks. **D**. CSPs and residues experiencing line broadening mapped onto the structure of WDR5. **E**. Binding curves of UMLILO to WDR5 in the presence of WDR5 inhibitors. Error bars represent s.e.m, n = 3. **F**. ^1^H-^15^N TROSY NMR spectra of ^15^N-labeled WDR5^23–334^ alone (black) and in the presence of an equimolar amount of UMLILO SL4 or a twofold molar excess of MM-102 or WDR5-IN6 (red). **G**. Linear map of WDR5 with residues exhibiting CSPs or line broadening indicated. SL = stem-loop, CSP = chemical shift perturbation, LB = Line broadening based on change (Δ) in peak intensity, µ = mean, 2σ = two standard deviations from the mean.

Titrating in RNA caused both chemical shift perturbations (CSPs) and line broadening of amide resonances clustered around blades 4 and 6 (**Fig. 5B–D, Supplemental Fig. 5A**). Many of the residues that experience CSPs were buried within the hydrophobic core away from the protein surface, suggesting that RNA binding imparts structural changes in WDR5. In agreement with this, the addition of RNA resulted in the increased intensity of amide resonances around the central core, base of blade 5, and residues in unstructured loops involved in MLL1 and H3K4 peptide binding (e.g., D107, L234, K259^36^; **Supplemental Fig. 5B**). We hypothesize that the presence of RNA may stabilize these flexible domains, but more work is required to confirm this mechanism. Overall, the clustering of spectral changes to at blades 4 and 6 is consistent with the literature: a large macrocyclic peptide spanning the cleft between blades 5 and 6 disrupts WDR5–HOTTIP lncRNA binding, and acylating amino acids on these binding surfaces results in reduced binding with UMLILO^37,38^.

WDR5 contains two known binding pockets: (1) the WIN-site, which mediates chromatin and MLL1 interactions, and (2) the WBM site, which binds MYC, the RbBP5 subunit of the MLL1/WDR5 complex, and the proposed lncRNA binding site^39,40^. Small-molecule inhibitors targeting the WIN and WBM binding pockets were used to determine whether UMLILO binding was disrupted by occupancy of these sites. We found that, in agreement with previous studies^5^, WDR5 binding to UMLILO was not inhibited by the WIN-site inhibitor MM-102 (DMSO control K_D_ = 52 ± 2 nM; MM-102 K_D_ = 40 ± 2 nM), but surprisingly was slightly enhanced (**Fig. 5E**). Targeting the WBM pocket with the small molecule WDR5-IN6 only partially disrupted WDR5–UMLILO binding (K_D_ = 73 ± 9 nM), which suggested that UMLILO binds WDR5 through contacts outside of the canonical WBM site.

To investigate whether the modest change in binding affinity was due to competition for binding site residues by the small molecules, we turned to NMR. The majority of spectral changes for both small molecules were distinct from those induced by UMLILO SL4 (**Fig. 5F, Supplemental Fig. 5C**). As expected, MM-102 caused CSPs and line broadening of residues at the WIN site (**Supplemental Fig. 5D**) and WDR5-IN6 showed CSPs of residues at the WBM site consistent with crystal structures^17,18^. MM-102 showed greater line broadening and CSPs overall than WDR5-IN6, supporting previous work that showed small molecules occupying the WIN site induce structural changes and reduce the dynamics of allosteric regions^22,41^. We suspect that MM-102–induced changes in WDR5 structure and dynamics are potentially how it subtly enhances RNA binding. WDR5-IN6 showed perturbations outside the WBM site, possibly due to transient binding to other aromatic regions or induced allosteric effects. Together, these data support the conclusion that lncRNAs bind to WDR5 across multiple binding surfaces, including residues surrounding the WBM pocket and on the top face near blade 4.

### UMLILO influences WDR5 condensate formation and morphology

We investigated whether UMLILO has an impact on WDR5 LLPS. WDR5 condensates form as a result of a self-interaction between the WIN-like motif at the N-terminus of the IDR and the WIN site and do not form when the IDR is truncated^8^ (**Fig. 6A**). UMLILO readily partitions into WDR5 condensates but is not sufficient to induce phase separation in the absence of the self-association of WDR5 (**Fig. 6A)**.

**Figure 6.**
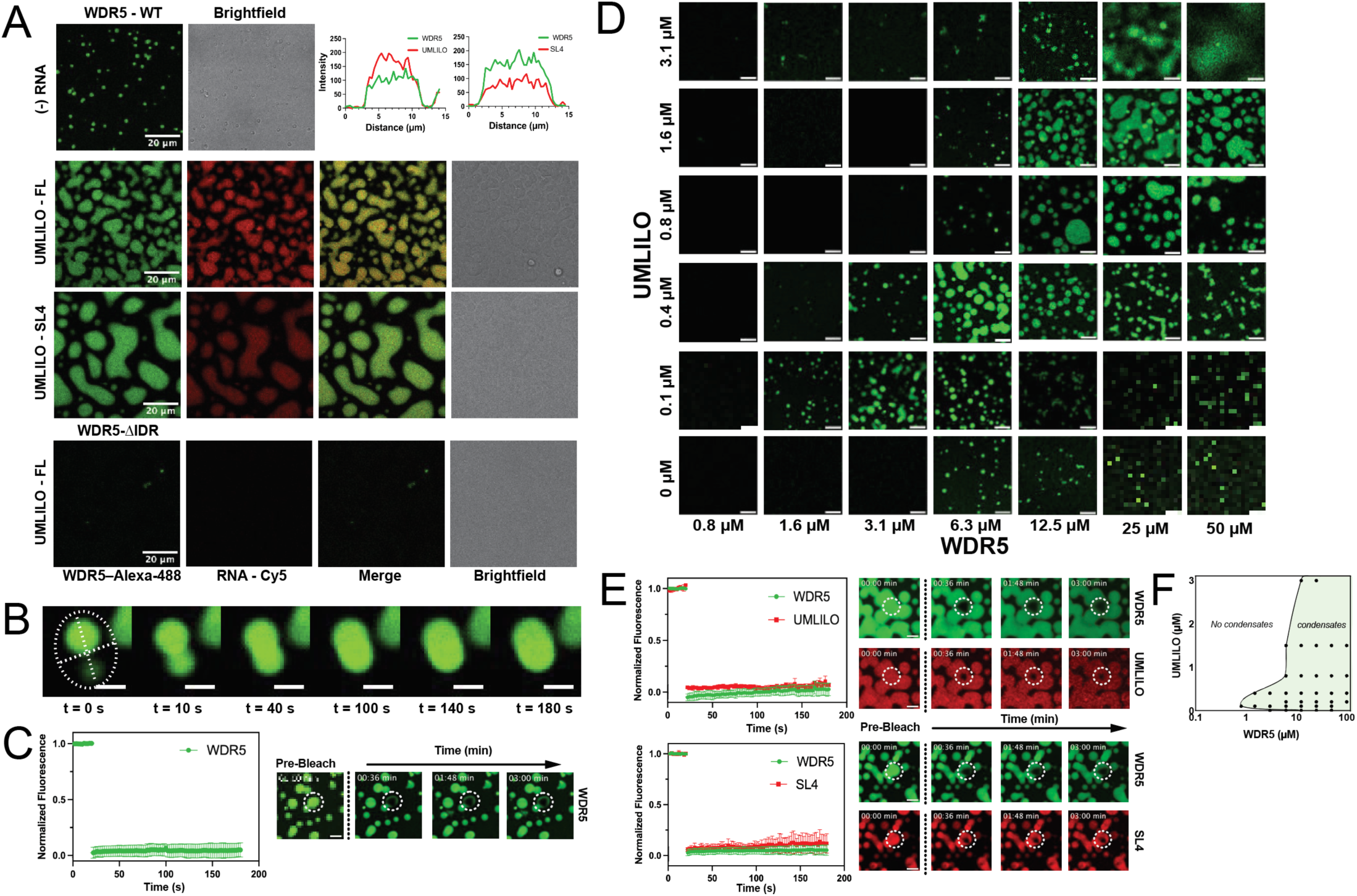
RNA increases phase separation propensity of WDR5 without changing dynamics. **A.** Composite micrographs of WDR5 and WDR5-ΔIDR labeled with Alexa-488 in the presence of UMLILO and UMLILO SL4 labeled with Cy5. Final protein concentrations were as follows: 60 µM WDR5 in the RNA-free sample, 13 µM WDR5 for samples with RNA, and 75 µM WDR5-ΔIDR. Final concentrations for UMLILO and SL4 were 400 nM and 5.75 µM, respectively. **Inset plots**: line profile inside condensate of fluorescence intensity of labeled WDR5 (green) and RNA (red). **B**. Coalescence event of WDR5 condensate over the course of 180 seconds. Scale bar = 2 µM. **C**. Fluorescence recovery after photobleaching (FRAP) analysis of WDR5-Alexa-488 in the absence of RNA. Scale bar = 5 µM, error bars represent s.e.m. **D**. Phase diagram of WDR5 in the presence of UMLILO. Scale bar = 10 µM; error bars represent s.e.m, n = 3. **E.** FRAP analysis of WDR5-Alexa-488 in the presence of RNA-Cy5. Scale bar = 10 µM; error bars represent s.e.m, n = 3 **F**. Graphical representation of WDR5–UMLILO phase diagram. ΔIDR = WDR5^23–334^ FL = full length (575 nt) UMLILO, SL4 = UMLILO stem-loop 4.

RNA identity influenced condensate formation and morphology (**Fig. 6A, Supplemental Fig. 6A**). WDR5 can undergo phase separation in the absence of RNA; however, co-incubation with UMLILO increased the overall phase separation propensity of the protein, resulting in an increase in the protein-dense phase (**Fig. 6A&D**). Interestingly, even though both full-length UMLILO and SL4 readily partitioned into WDR5 condensates, addition of SL4 significantly increased the size and wetting of the condensates (**Fig. 6A&D).** Condensate formation, shape, and size also changed as a function of RNA concentration, with phase separation promoted at low nanomolar concentrations of UMLILO and disrupted when UMLILO concentrations exceeded an approximately 1:4 RNA:protein ratio (**Fig. 6D**).

We next sought to investigate the internal dynamics of the condensates, and the potential impact the distinct RNA identities might have on these dynamics, by performing fluorescence recovery after photobleaching (FRAP) experiments. Under the conditions used in this study, there was no recovery of either fluorescently labeled WDR5 or UMLILO. This suggests that although the RNA partitions within the condensates, there is no free diffusion of either of the macromolecules. The internal dynamics of condensates were unaffected by the addition of a crowding agent and linear 30 nt-poly(U) RNA (**Supplemental Fig. 6B&C**). Interestingly, the condensates were capable of relaxing to undergo fusion as a function of time, even in the absence of FRAP recovery (**Fig. 6B),** suggesting the condensates have a viscoelastic state dictated for the most part by the WDR5 network. These results suggest that although the condensate morphology might be affected by the overall RNA concentration and identity, WDR5 self-interactions dictate the condensate’s overall material state.

## Discussion

The regulatory mechanisms that underlie trained immunity are complex, involving networks of RNA–DNA, RNA–protein, and protein–protein interactions. In this study, we have investigated the role of UMLILO and its interaction with WDR5 to stimulate gene expression in the CXCL TAD. UMLILO is rapidly transcribed upon innate immune cell activation by an environmental stimulus, and the high affinity of UMLILO for WDR5 enables its recruitment to the specific genomic locus.

Our work reveals that WDR5 interacts with the stem-loops of UMLILO, and this multivalency increases overall affinity. A bipartite binding mechanism with highly structured loops confers specificity, while flexible single-stranded regions provide an electrostatic velcro-like effect that can stabilize the complex. WDR5 does not promiscuously bind with low nanomolar affinity to any RNA but can interact transiently with mid-nanomolar affinity to other structured RNAs and has the potential to bind tightly to G-quadruplex-containing lncRNAs. These multiple binding modes, linked to RNA structural properties, could explain how WDR5 interacts with other RNAs across different contexts, with RNA structure dictating the nature of the interaction^27,28,42^. In agreement with this, the sequence and structural conservation of UMLILO supports the idea that WDR5 recognizes specific elements.

Although several studies have investigated how WDR5 binds peptides and small-molecule inhibitors, little *in vitro* work has been carried out to investigate how WDR5 recognizes and binds RNA. It was previously reported that a phenylalanine at position 266 was primarily responsible for RNA binding^5^. Our extensive *in vitro* characterization of the WDR5–UMLILO interaction shows that F266 is essential for WDR5 stability at physiological temperatures but not required for UMLILO binding. This finding is important for contextualizing work in which F266A is used as an *in vivo* control to abrogate lncRNA binding^28,31,38^, as not all lncRNA interactions may be disrupted. Further *in vitro* binding assays between the F266A mutant and HOTTIP, as well as additional *in vivo* validation, are required to confidently attribute cellular phenotypes of the F266A mutant to loss of lncRNA binding.

We found that WDR5 binds RNA at multiple surface sites, consistent with the many aromatic and polar RNA-interacting residues present in WD40 repeats. We conclude that the two binding surfaces enable the multiple binding modes of WDR5 and explain why it interacts differently with each stem-loop. These binding sites overlap but are distinct from the WIN and WBM binding sites, which explains how lncRNAs and other members of the WDR5/MLL1 complex, such as RBBP5, can bind WDR5 simultaneously^5^. Interestingly, RNA binding appears to influence WDR5 dynamics, increasing the rigidity of amino acids in the loops near the MLL1 and histone H3 binding site responsible for ligand recognition and orientation^36,43,44^. This may entropically favor MLL1 binding or position the loops to create a binding surface favorable for MLL1 binding, ultimately improving recruitment and assembly of the WDR5/MLL1 complex.

The presence of multiple RNA-binding sites creates the potential for more transient interactions to enhance phase separation. WDR5 has been demonstrated to partition into biomolecular condensates^7,8^. Understanding the role of this biophysical phenomenon in the context of transcription regulation is in its infancy; however, our results, combined with the research carried out by others, establish a foundation for understanding this process. Rapid transcription of UMLILO has the potential to drive phase separation within the *CXCL* chemokine TAD, thereby enriching for WDR5 and preventing its diffusion away from gene promoters. In conjunction with this, it has been shown that WDR5 and MLL1 can partition together in condensates, with MLL1 possessing increased catalytic activity when WDR5 is present^6^. Together, the enhanced phase separation of WDR5 induced by UMLILO lends support to its super-enhancing properties, as phase separation is often a mechanistic explanation for why lncRNAs have an outsized effect relative to their transcript abundance.

Based on our results, together with findings from others, we propose a hypothetical model in which UMLILO initiates phase separation by bringing multiple WDR5 molecules into close proximity. After extensive transcription, excess UMLILO disrupts the phase separation within the CXCL TAD, thereby releasing WDR5 back into the nucleus. In addition to excess RNA disrupting condensate formation, at high concentrations RNA may disrupt RBBP5 or MLL1 binding because their binding pockets partially overlap. Furthermore, we suspect that UMLILO may act as a scaffold, tethering WDR5 to histone proteins and thereby enhancing localization and residence time near the methylation site. Future work investigating this in cells is a promising direction towards a full understanding of the structure-function roles of UMLILO and WDR5 in regulating gene expression.

## Methods

### In vitro transcription and purification of RNA

The DNA template encoding for UMLILO was amplified by polymerase chain reaction (PCR) from a plasmid with the exonic sequence (a gift from Dr. Musa Mhlanga) using a forward primer containing the T7 promoter sequence. DNA templates encoding shorter constructs and T7 promoter sequence were obtained from Integrated DNA Technologies (IDT). The transcription reaction consisted of 0.64 µM DNA template, 8 mM of each rNTP, 10% polyethylene glycol 8000, 20-40 mM MgCl_2_, 1× transcription buffer (5 mM Tris, pH 8.5 mM spermidine, and 10 mM dithiothreitol (DTT)), and in-house expressed and purified T7 bacterial RNA polymerase. RNA was *in vitro* transcribed at 37°C for 3 hours, then purified by high-performance liquid chromatography (HPLC) on a DNAPac^TM^ PA200 column (Thermo Scientific) using 12.5 mM Tris HCl and 6 M urea and eluted with a gradient of 0-500 mM sodium perchlorate. RNA was desalted using Sep-Pak® Plus Long C18 columns (Waters) or dialysis, then lyophilized and resuspended in water.

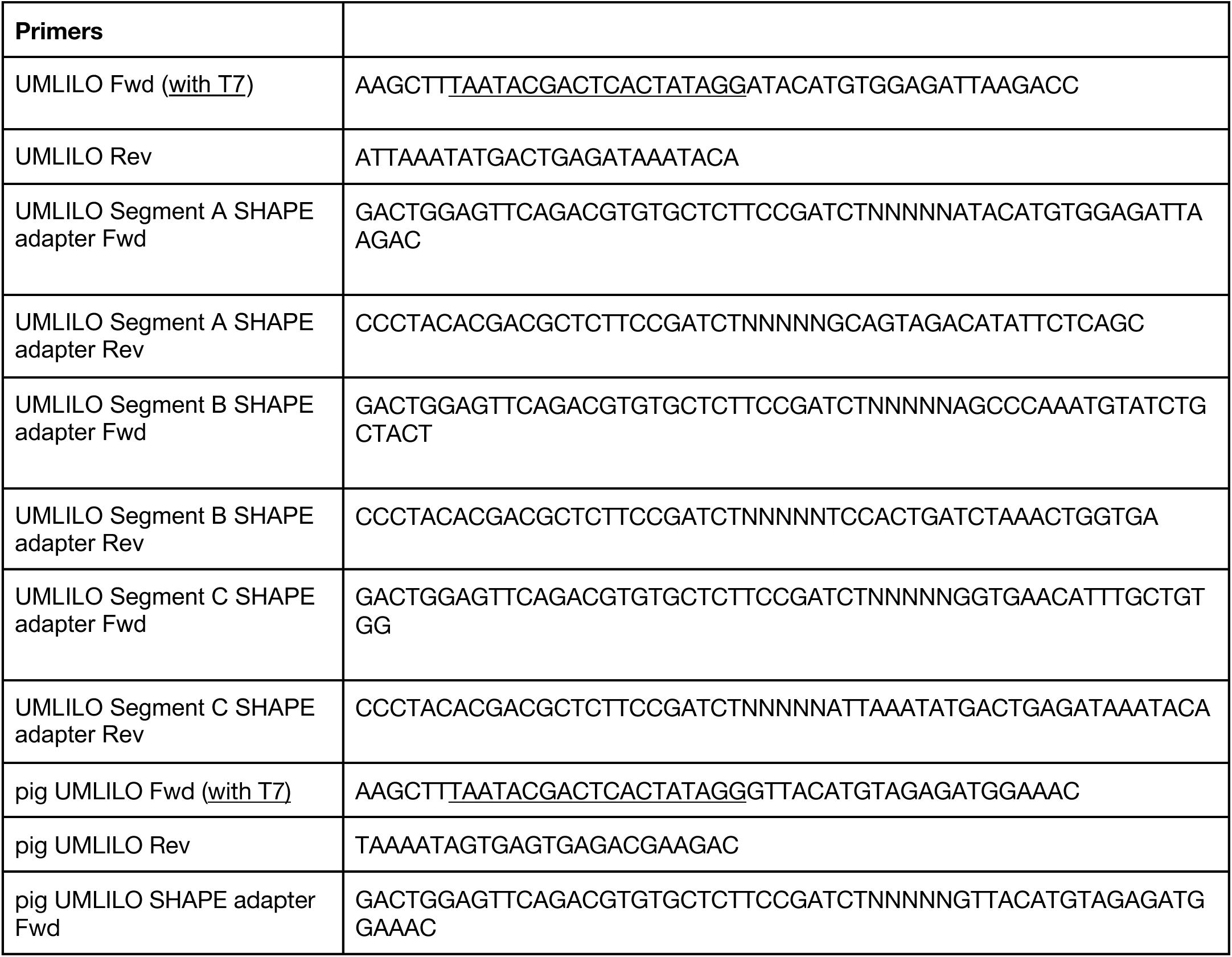
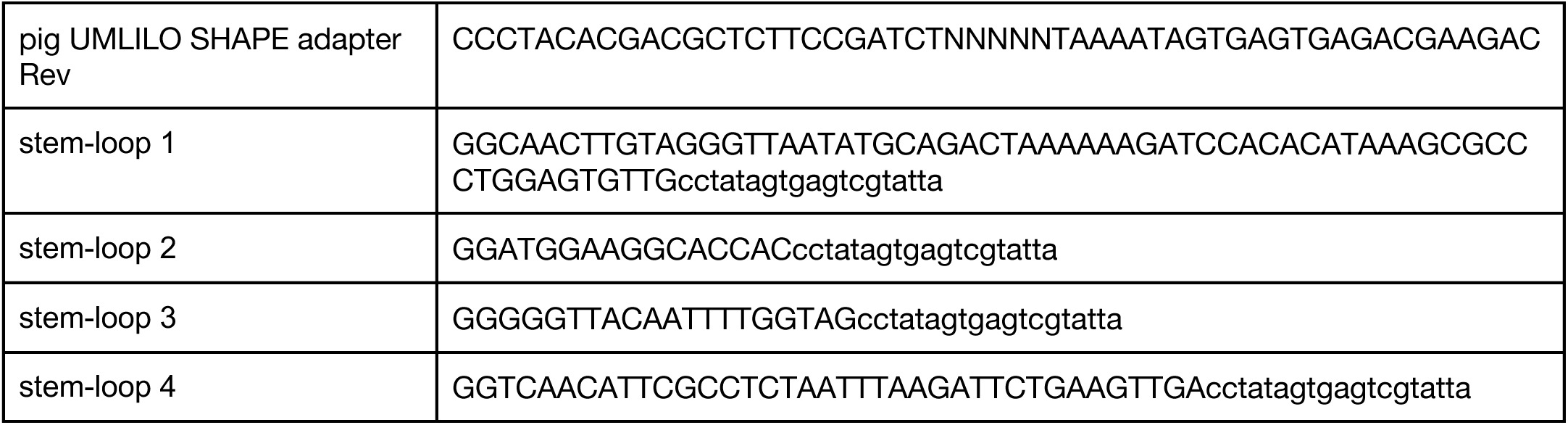

### Expression and purification of WDR5

Plasmid pGEX-4T1-WDR5 was obtained from Addgene (Addgene plasmid # 59969)^45^. The sequence encoding for WDR5 was amplified by PCR and ligated into pETM11 with an N-terminal 6x-His tag and Tobacco Etch Virus (TEV) cleavage site. The WDR5^23–334^ construct was made by Gibson Assembly. A gBlock encoding for the F266A mutant was ordered from IDT and ligated into pETM11. Plasmids were each transformed into BL21 DE3 *E. coli* competent cells. LB broth supplemented with 50 µg/mL kanamycin was inoculated with an overnight culture and incubated at 37°C until the optical density (OD_600_) reached 0.8-1.2. Expression was induced with 1 mM isopropyl β-D-1-thiogalactopyranoside (IPTG) at 18°C overnight. Cells were lysed by sonication in a buffer containing 50 mM HEPES, pH 7.5, 300 mM NaCl, and 1 mM DTT then purified by immobilized metal affinity chromatography (IMAC) (NEBExpress^®^ Ni Resin S1428S) and eluted with 100 mM imidazole, followed by dialysis and cleavage with TEV protease made in-house. The protein was further purified by size exclusion chromatography on a Superdex S75 column in a buffer containing 50 mM HEPES, pH 7.5, 150 mM NaCl, and 1 mM DTT. Fractions were analyzed by SDS-PAGE, flash-frozen, and stored at -80°C.

^15^N-labeled WDR5^23–334^ was prepared in minimal media containing trace elements (134 µM ethylenediaminetetraacetic acid (EDTA), 3.1 µM FeCl_3_, 0.62 µM ZnCl_2_, 76 nM CuCl_2_, 42 nM CoCl_2_, 162 µM H_3_BO_3_, 8.1 µM MnCl_2_), M9 salt solution (33.7 mM Na_2_HPO_4_, 22 mM KH_2_PO_4_, 8.55 mM NaCl, 9.35 mM ^15^NH_4_Cl), 0.3 mM CaCl_2_, 1 mM MgSO_4_, 0.4% glucose, 1 µg/mL biotin, 1 µg/mL thiamin, and 50 µg/mL kanamycin. The culture was incubated at 37°C until OD_600_ reached 0.75, induced with 1 mM IPTG at 18°C, incubated at 18°C overnight, then purified as before. Size-exclusion chromatography was performed in an NMR buffer containing 25 mM sodium phosphate, pH 6.0, 50 mM NaCl, and 1 mM DTT. Samples were stored at 4°C for up to 3 weeks to prevent precipitation upon thawing, by which time no degradation (as assessed by SDS-PAGE) nor loss of binding was observed.

### Filter dot blot assays

Filter dot blot binding assays were performed in a buffer of 25 mM HEPES, pH 7.5, 50 mM NaCl, and 0.5 mM DTT unless noted otherwise. 3′ Cy5-labeled RNA (6 nM final concentration) was combined with a dilution series of WDR5 in a total volume of 55 µL, incubated on ice for 30 min, then applied to a filter dot blot apparatus with nitrocellulose and nylon membranes (pore sizes 0.45 µm) and gently drawn through by vacuum. For filter dot blots with WDR5-IN6 (MedChemExpress HY-154949) and MM-102 (Sigma-Aldrich 5006490001), WDR5 was incubated with WDR5-IN6 (75 µM), MM-102 (300 µM), or an equivalent volume of DMSO for 15 min at room temperature, then mixed with RNA, and blots carried out as before. Membranes were imaged using a Bio-Rad Chemidoc imaging system and quantified in Fiji ImageJ^62^. All assays were carried out in technical triplicate, binding curves were fitted to the Hill equation and plotted in GraphPad Prism 10, ± denotes standard error of the mean (s.e.m.).

### Circular Dichroism (CD) and Thermal Denaturation Assays

CD spectra were collected using a Jasco J-1500 Circular Dichroism spectrometer with wavelengths scanning from 190 nm to 300 nm. Spectra were recorded using a quartz cuvette with a 1 mm path length, with a protein concentration of 10-15 µM in a buffer containing 50 mM HEPES, pH 7.5, 50 mM NaCl, and 1 mM DTT. Scans were performed at 25°C, and each final spectrum was the average of three accumulation trials collected with a step size of 1 nm a time per point of 1 s and a bandwidth of 1 nm. A buffer-only blank was subtracted from each spectrum. Thermal denaturation assays were performed at 213 nm, starting at 25°C and increasing to 95°C at 1°C/min. Spectra were collected at 1°C intervals.

### Chemical probing

Chemical probing was performed as described previously^46–48^. Briefly, *in vitro* transcribed RNA was heated to 95°C for 5 min, then immediately cooled on ice. RNA was refolded at 37°C in a buffer containing 100 mM HEPES, pH 8, 100 mM NaCl, and 10 mM MgCl_2_. For all chemical probing experiments, multiple probe concentrations were screened to ensure single-hit kinetics. For SHAPE-MaP experiments, RNA was incubated with 6 mM 1-methyl-7-nitroisatoic anhydride (1M7, synthesized in house) for 7 minutes at 37°C, and then quenched with 333 mM DTT. For DMS-MaP experiments, RNA was probed with 10.1 mM dimethyl sulfate (DMS) at 37°C for 15 min and quenched with 333 mM DTT. The RNA was cleaned up with RNAClean XP beads, and SHAPE samples were reverse transcribed using SuperScript II™ (Invitrogen™, 18064022), while DMS samples were reverse transcribed using Induro Reverse Transcriptase (New England Biolabs, M0681L). The resulting cDNA was purified using RNAClean XP beads, amplified with Q5 Hot Start (NEB, M0491S), and PCR products cleaned using AMPureXP beads (Beckman, A63881). Sequence specific primers and Illumina adapters were ordered from IDT. 1M7-probed RNA library quality was checked using an Agilent 4200 TapeStation and sequenced on an Illumina MiSeq system with MiSeq Reagent Kit v3. DMS-probed RNA was sequenced using Oxford Nanopore Technologies Flongle Flow Cell (ONT, FLO-FLG114) with MinKNOW v25.09.16 software. Chemical probing in the presence of WDR5 was performed on refolded RNA cooled to room temperature in a buffer containing 50 mM HEPES, pH 7.5, and 100 mM NaCl. Samples were then incubated with either 40 mM 1M7, 6 mM 5-nitroisatoic anhydride (5-NIA), or 20.2 mM DMS at room temperature for 15 min. RNA cleanup and reverse transcription were performed as before. The resulting cDNA was amplified using PrimeSTAR GXL DNA polymerase (Takara Bio, R050A), cleaned as before, then sequenced by Oxford Nanopore Technologies as before. Chemical probing of pig UMLILO was performed following the same protocol; DMS treated samples were sequenced on an Illumina MiSeq system, and 5-NIA treated samples sequenced by Oxford Nanopore as before. Sequencing results were analyzed using ShapeMapper2, deltaSHAPE, RNA Framework, and visualized using RNAcanvas^47,49–51^. DaVinci analysis was done with 5-NIA probed RNA sequenced by Oxford Nanopore^35^.

### Motif analysis

Alignment of UMLILO across 43 species was performed using sequences extracted from Multiz DNA alignment available from the UCSC Genome Browser (http://genome.ucsc.edu, GRCh38/hg38 assembly) and aligned to the human 575-nt transcript using Clustal Omega. PhyloP and PhyloCons base-wise conservation scores were obtained from the Hiller Lab 470 Mammals genome alignment obtained from the UCSC Genome Browser^52,53^. The sequence logos were created using WebLogo 3^54^. R-scape was used to identify conserved structural elements in the alignment, using a windowed approach shown to be more accurate for lncRNAs^55,56^. A window size of 130 nts, sliding by 30 nts, was sufficient to capture full secondary structural elements and identify conserved nucleotides with an E-value cutoff of 0.05. LocARNA multiple alignment of UMLILO domain II was used with the Local alignment type option^57–59^.

### Nuclear magnetic resonance (NMR) spectroscopy

All NMR experiments were carried out on a Bruker AVANCE III 600 MHz NMR equipped with a 5 mm TCI CryoProbe. For 1D ^1^H imino proton titration experiments, WDR5^23–334^ was titrated into 90 µM SL4 in a buffer containing 25 mM sodium phosphate, pH 6.0, 50 mM NaCl, and 1 mM DTT, supplemented with 10% D_2_O. For ^1^H-^15^N BEST-TROSY titration experiments, unlabeled UMLILO SL4 was titrated into ^15^N-labeled WDR5^23–334^ diluted to 90-110 µM in a buffer containing 25 mM sodium phosphate, pH 6.0, 50 mM NaCl, and 1 mM DTT, supplemented with 10% D_2_O. WDR5-IN6 and MM-102 were dissolved in neat DMSO, and titrations were performed following the same experimental methods. All experiments were performed at 298 K. Data were processed in TopSpin (Bruker RRID: SCR_014227) and analyzed using CcpNmr Analysis Version 3^60^. NMR assignments were obtained from BMRB entries 27528^61^ and 52291^7^.

### Condensate Experiments

WDR5 constructs were labeled with Alexa Fluor™ 488 NHS Ester (Invitrogen™, A20000) and RNA was 3′ Cy5 labeled (Jena Bioscience, NU-1706-CY5). Coverslips with custom-made wells of approximately 6 µL in volume were used for incubation and imaging. The condensate buffer was composed of 50 mM HEPES, pH 7.5, 50 mM NaCl, and 1 mM DTT and 0.5-1% labeled species were used for each sample. Condensates were induced by diluting WDR5 to its desired final concentration along with RNA. Samples were incubated for 1 h at room temperature prior to imaging. Confocal images were taken either on a Leica TCS SP8 X Laser Confocal Microscope equipped with a 63×/1.30 NA PLAN APO objective, using a white-light laser source and a Leica DFC310 digital color camera or a Zeiss LSM800 laser scanning confocal microscope equipped with two GaAsP detectors (Carl Zeiss, Oberkochen, Germany) using a 63×/1.4 NA Plan-Apochromat oil objective with excitation at 488 nm, and a pinhole set to 1 Airy unit. Images were processed and analyzed using Fiji ImageJ.

For FRAP experiments, single-slice confocal images were acquired using a Marianas Spinning Disk confocal microscope (Intelligent Imaging Innovations) consisting of a spinning disk confocal head (CSU-X1; Yokagawa) on a Zeiss Axio Observer inverted microscope equipped with ×63/1.4-NA Plan-Apochromat (oil immersion). Images were acquired with a Prime sCMOS camera (Photometrics) controlled by SlideBook 6 (Intelligent Imaging Innovations). Alexa Fluor™ 488 or Cy5 were excited with the 488-nm and 637-nm lines, respectively, from a solid-state laser (LaserStack) and collected with a 440-/521-/607-/700-nm quad-emission dichroic and 525-/30-nm or 692/40-nm emission filter. Regions of interest with diameters of 0.5 µm were bleached with a 488 nm laser line. Fluorescence recovery was monitored every 2 seconds for 90 time points. Condensates were induced at a final concentration of 12.5 µM WDR5 with either no RNA, 0.4 µM full-length UMLILO RNA, 5.75 µM SL4 RNA, or 5 µM 30-nt poly(U).

## Data Availability

Sequencing data is available at NIH BioProject PRJNA1502428.

## Acknowledgements

The authors acknowledge the NYU Department of Chemistry Shared Instrumentation Facility, the shared facilities provided through the Materials Research Science & Engineering Center (MRSEC) program of the NSF under Award Number DMR-1420073, and the NYU Langone Microscopy Core which is partially supported by NYU Cancer Center Support Grant NCI P30CA016087. We thank the lab of Evgeny Nudler for assistance with Illumina sequencing. We thank Seungwoo Lee for assistance with NMR experiments, Harsha Venkatesh for assistance with DaVinci analysis and circular dichroism spectroscopy, Shengwei Xiong for synthesizing 1M7, and the members of the Jones lab for their insights and feedback. A.N.J. is supported by NSF 224667 and NIH 1R35GM160278. A.C. is supported by NIH T32GM158479 and the Margaret Strauss Kramer Fellowship.

## Author Contributions

A.C. and A.N.J. conceptualized the project, A.C. and A.V.C. carried out experiments and analyzed results, A.C. and A.V.C. made the figures, A.C. and A.N.J. drafted the manuscript, and A.C., A.V.C., and A.N.J. revised the manuscript. A.C. and A.N.J. secured funding. A.N.J. supervised the project.

