## Supplement for "Conserved RNA structural elements mediate WDR5 recognition of the long non-coding RNA UMLILO"

### Supplemental Information

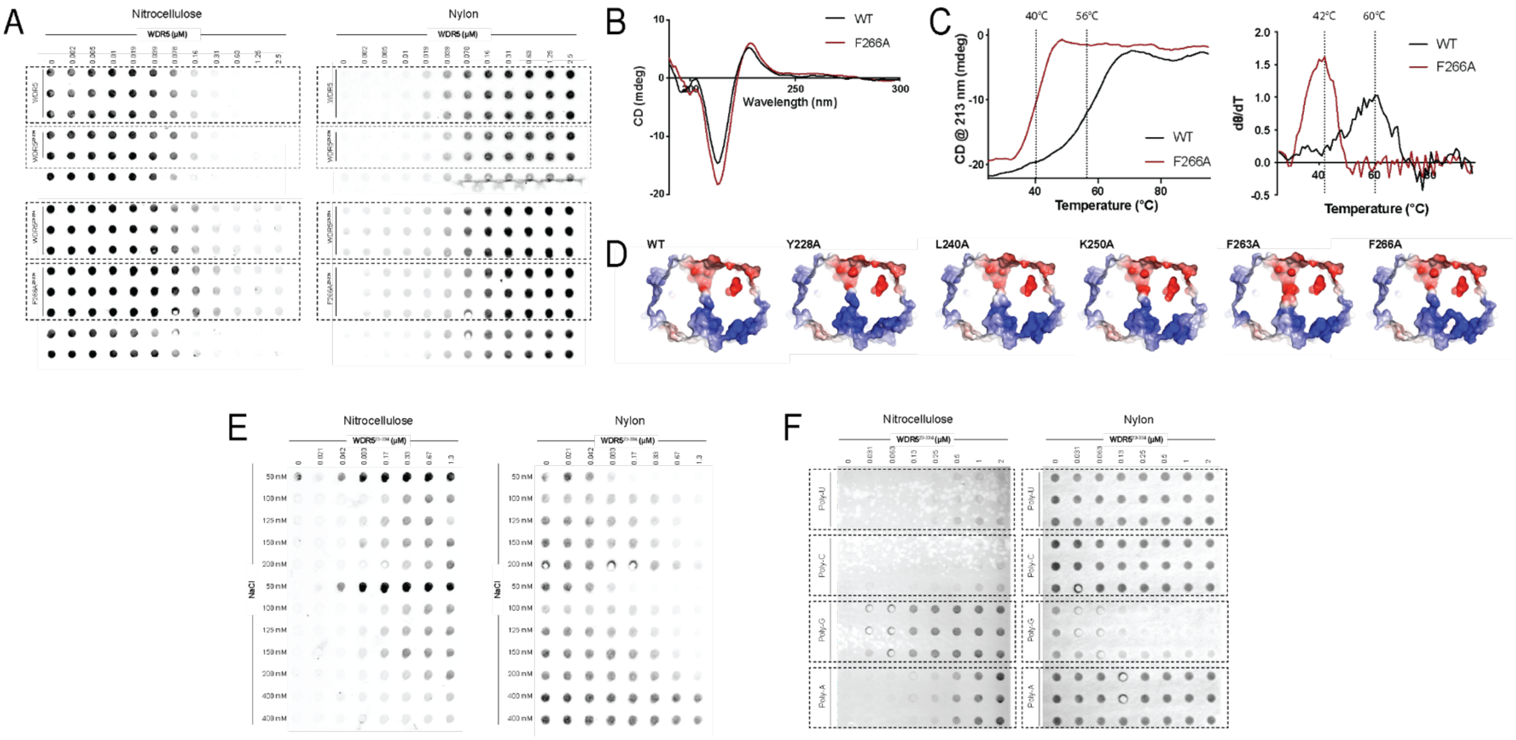


**Supplemental Figure 1. A.** Filter dot blot assays of UMLILO and each WDR5 construct. **B.** Circular dichroism (CD) spectra for wild type WDR5 and F266A mutant. **C.** CD thermal melting curve and its first derivative collected at 213 nm for wild type WDR5 and F266A mutant. **D.** Cross sections of AlphaFold 3 models of $WDR5^{30\text{-}334}$ point mutants. **E.** Filter dot blot assays of UMLILO and WDR5 under increasing ionic conditions. **F.** Filter dot blot assays of WDR5 binding to poly(A), poly(C), poly(G), and poly(U) RNAs.

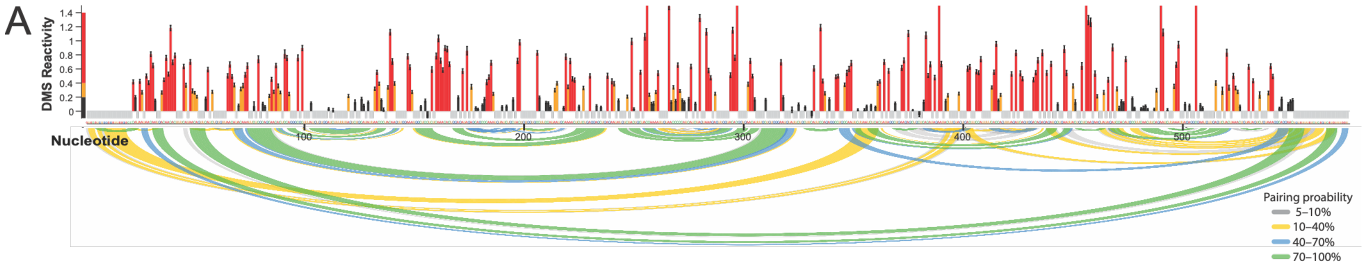


**Supplemental Figure 2. A.** DMS reactivity profile (top) and pairing probabilities for UMLILO (bottom).

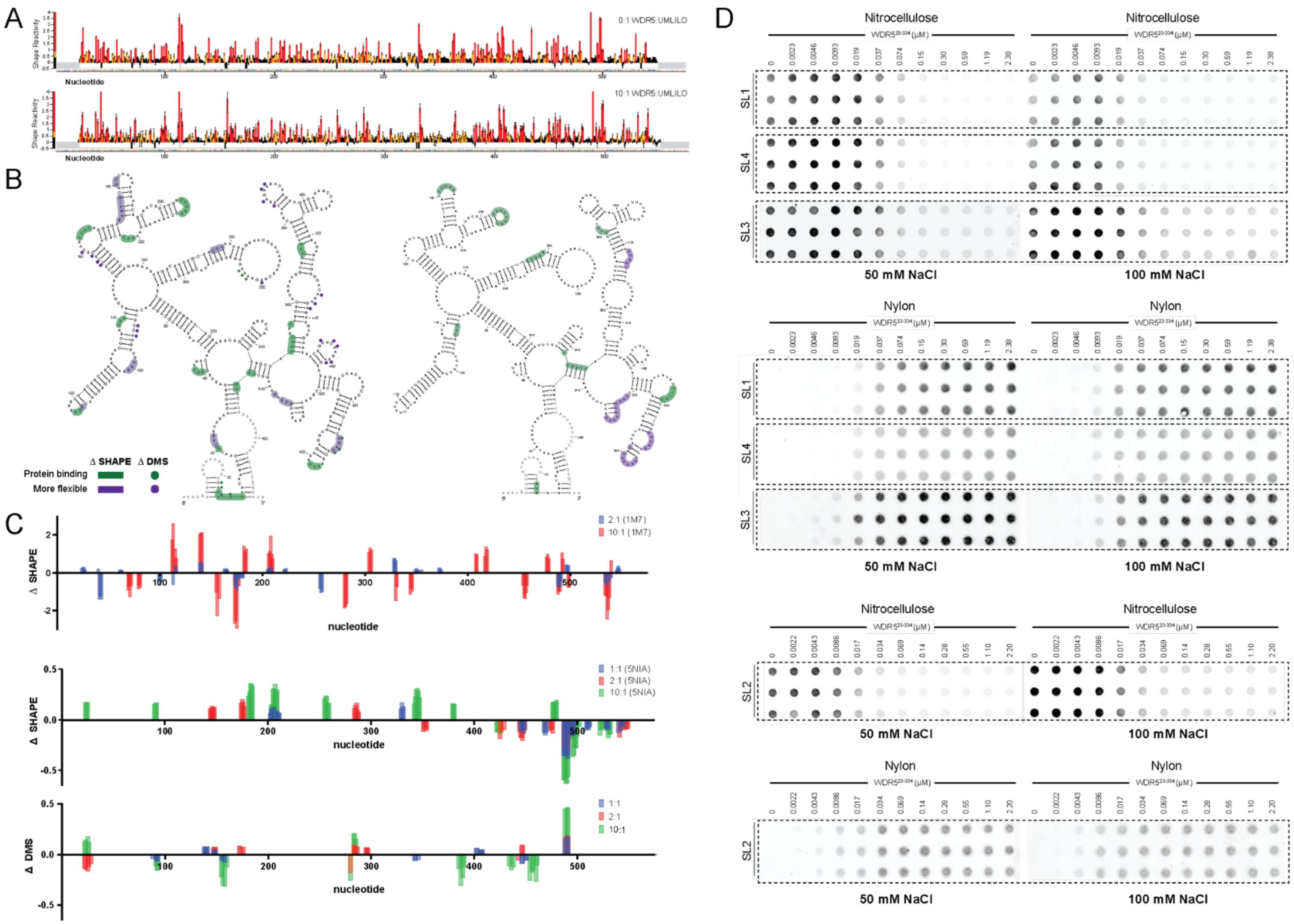


**Supplemental Figure 3 A.** 1M7 reactivity profiles of UMLILO with and without WDR5. **B.** Changes in reactivity to 1M7 at 2:1 ratio of WDR5 to UMLILO (left) and to 5NIA at 10:1 ratio of WDR5 to UMLILO (right). **C.** ΔSHAPE and ΔDMS plots for different ratios of WDR5 and UMLILO with 1M7, 5-NIA, and DMS chemical probes. **D.** Filter dot blots of each stem loop and WDR5$^{23\text{-}334}$.

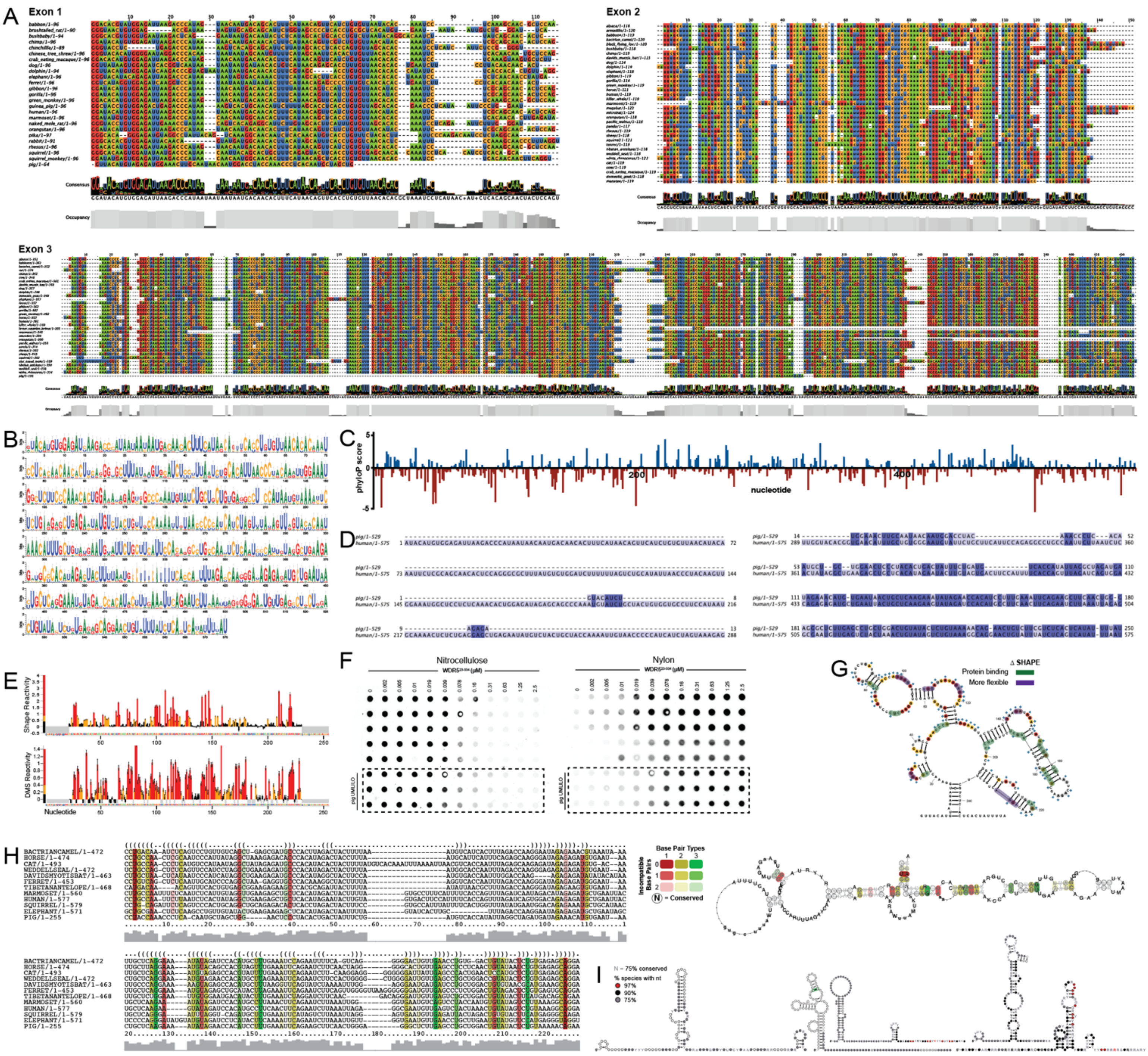

**Supplemental Figure 4. A.** Sequence alignment of each exon, colored by nucleotide identity. **B.** WebLogo generated from UMLILO sequence alignment. **C.** Plot of phyloP conservation scores from Hiller 470-way mammal alignment. A positive score indicates evolutionary conservation of the nucleotide while a negative score indicates accelerated evolution. **D.** Pairwise sequence alignment of human and pig UMLILO colored by nucleotide similarity. **E**. 5-NIA and DMS reactivity profiles for pig UMLILO. **F.** Filter dot blot binding assays of pig UMLILO and WDR5. **G**. Changes in reactivity to 5-NIA and DMS mapped on pig UMLILO secondary structure. **H**. LocARNA multiple alignment and consensus structure for the entirety of domain II of UMLILO. **I**. R-Scape results for each 130 nt window containing major structural elements of UMLILO.

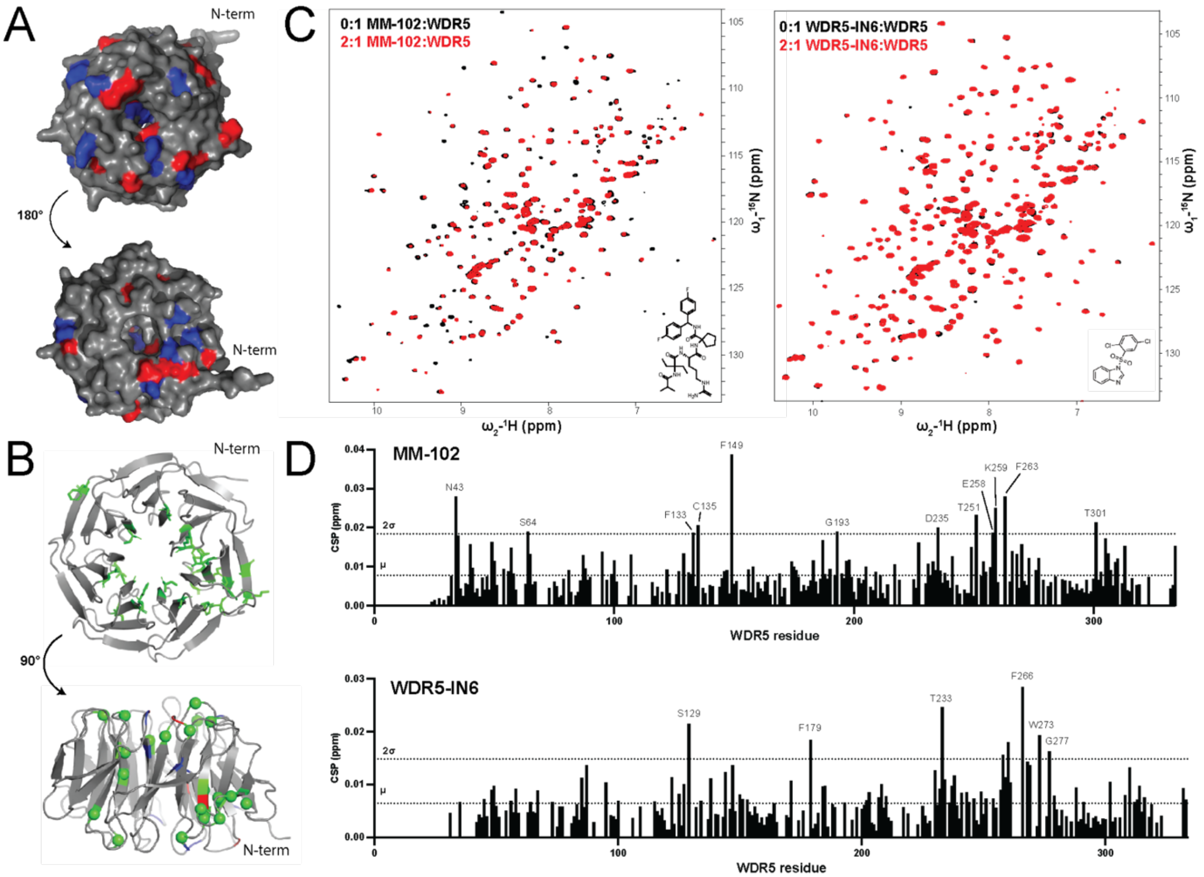


**Supplemental Figure 5. A.** WDR5 surface mapped with chemical shift perturbations (blue) and line broadening (red) one standard deviation away from the mean. **B.** Top and side views of WDR5 with residues that increased in peak intensity upon RNA titration into WDR5 colored in green. **C**. TROSY spectra for titrations of MM-102 and WDR5-IN6 into WDR5. **D.** CSP plots for MM-102 and WDR5-IN6. μ = mean, 2σ = 2 std deviations.

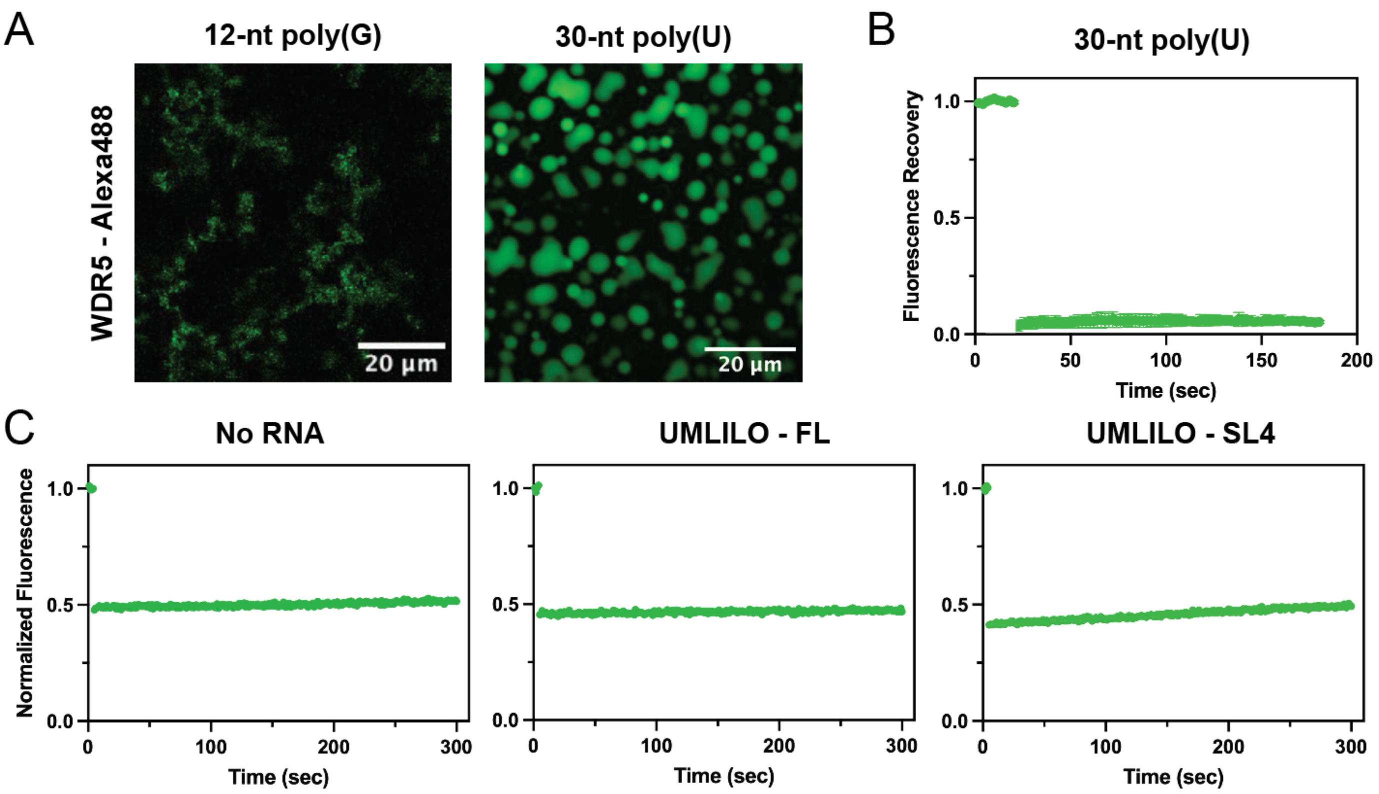


**Supplemental Figure 6. A.** Confocal images of WDR5 in the presence of 12-nt poly(G) and 30-nt poly(U) RNAs at 2 and 5 µM, respectively. Protein concentration was 13 µM. **B.** Normalized fluorescence recovery of WDR5 in the presence of 30-nt poly(U) RNA. **C**. Normalized fluorescence recovery of WDR5 in the absence of RNA, WDR5 in the presence of UMLILO, or WDR5 in the presence of SL4 when in a buffer containing 25 mM HEPES, pH 7.5, 150 mM KCl, 1 mM DTT, and 10% PEG-8000. All protein was labeled with Alexa-488. SL = stem-loop

| RNA | Class/Role | Binds WDR5 | Motif location | YUUVAANU motif with flanking nts |
|---|---|---|---|---|
| UMLILO (x2) | IPL[3] | Yes | nts 478-485; 493-500 | CAUGCCUUCAACUUCAGAA UCUUAAAUUAGAGG |
| Pig UMLILO (x2) | IPL | Yes | nts 159-166; 174-181 | CAUCCUUUGAAAUUCAGAA GCUUCAACUGGGAG |
| AC063976 | IPL | Predicted | nts 889-896 | UCACAGAAAAUUUAAACUU GGAAUAUUUU |
| AMANZI[63] | IPL | Predicted | nts 774-781 | CAUAAUAACCUUUAAAGUA GAUAAUAUU |
| IPL-CCL2 | IPL | Predicted | nts 493-500 | AUGGUGGGACUUGAACU GCCUCAUGCA |
| LINC00880_1 (x4) | IPL | Predicted | nts 105-112; 699-706; 753-760; 846-853 | GCACCAUGACCUUGAACUU UCCAGCCUU |
| RP11_215E13 (x2) | IPL | Predicted | nts 852-859; | CAGACCAUGGUUUGAAGUC |

| | | | | |
|---|---|---|---|---|
| | | | 990-997 | UGGAUCUUC |
| RP11-35612 (x2) | IPL | Predicted | nts 182-189; 578-585 | CCAAAAUGGGUUUAAAGUCUCAGCCAUA |
| TCONS_00013373_2 | IPL | Predicted | nts 662-669 | AAGUGAGCUCUUUAAAGUGGCUAAAGUG |
| HOTTIP (x2) | gene priming (H3K4me3)[29] | Yes | nts1561-1568; 2948-2955 | UCUCUCUCUUUUUAAAAUUCUCGCCUCU |
| SRA (x2) | gene priming (H3K4me3) among other functions[42] | Yes | nts 358-365; 904-911 | AGCUUCUUUGUUUGAAUUUGCUCCUGAG |
| NEAT1_1 (x3) | gene priming (H3K4me3) among other functions[64] | Yes | nts 1374-1381; 1999-2006; 3019-3026 | GAGGCUACCAUUUAAAGUUUGCAGAUGA |
| NeST | gene priming (H3K4me3)[26] | Yes | nts 914-921 | GGCUUCUCUCUUUCAAAUGGAAGCTAGC |
| *Truncated (UUVAANU)* | | | | |
| AC079922-001 | IPL | Predicted | nts 531-537 | CUGCUUCGGAUUCAAAUUCUGACUCCUU |
| RP11-499E18 (x3) | IPL | Predicted | nts 240-246; 344-350; 644-650 | GCCUCCCGGGUUCAAGUGAUUCUCCUG |

**Supplemental Table 1. WDR5-binding lncRNAs containing 5′ YUUVAANU 3′ motif with 10 flanking nucleotides.** In the case of multiple instances, only the first is shown. Note that the motif is not present in the following RNAs, which have not been shown to bind WDR5 for H3K4me3 priming: MEG3, SLNCR1, LINC-PINT, PCAT2, PINK1 mRNA, circARID1A. Only one of the known IPLs, ILF3-AS1-001, did not contain the motif. Abbreviations: Y = C or U, V = A, C, or G, N = any nucleotide; AMANZI = A MAster Non-coding RNA antagoniZing Inflammation, HOTTIP = *HOXA* transcript at the distal tip, SRA = Steroid Receptor RNA Activator, NeST = *ne*ttoie *S*almonella pas *T*heiler's [cleanup Salmonella not Theiler's], SLNCR1 = SRA-like non-coding RNA, LINC-PINT = Long Intergenic Non-protein Coding RNA, P53 INduced Transcript, PCAT = Prostate cancer associated transcript 2, PINK1 = PTEN-induced kinase 1
